# Single-probe, amplification-free multiplex mRNA PNA-FISH maps bacterial gene expression in intact three-dimensional biofilms

**DOI:** 10.64898/2026.10.29.762896

**Authors:** Ana Barbosa, Heidi J. Smith, Albert Parker, Christopher J. Jones, Darla M. Goeres, Nuno F. Azevedo, Laura Cerqueira

## Abstract

Biofilms are spatially structured communities in which local microenvironments drive physiological differentiation, yet mapping gene expression inside intact three-dimensional biofilms remains challenging. Bacterial spatial transcriptomics methods typically rely on tiled probe sets, sequential hybridization rounds, or signal amplification. Here, we show that a single short, directly labelled peptide nucleic acid (PNA) probe per target is sufficient to quantify bacterial mRNA *in situ* without amplification. With all probes designed for a common hybridization condition, a transcript panel is resolved in one hybridization on a standard confocal microscope, in unmodified cells while preserving biofilm architecture. Benchmarked against the biphasic cycle of *Legionella pneumophila*, multiplex mRNA PNA-FISH recovers the expected transcriptional switch and agrees quantitatively with RT-qPCR in planktonic and biofilm populations. In flow-cell biofilms, it shows that sessile populations do not recapitulate the planktonic program and that transcriptional differences follow cellular aggregation state rather than depth.

## Introduction

Cellular specialization is a hallmark of microbial life, shaping the behavior and fate of individual cells within complex communities. Microbial life rarely exists in isolation but instead assembles into structured biofilms, where three-dimensional architectures generate spatially localized microenvironments that govern cellular physiology ^1–3^. The structural protection and metabolic cooperation of biofilms confer survival advantages inaccessible to planktonic cells, while steep physicochemical gradients drive genetically identical cells into distinct physiological states, producing stratified populations optimized for collective fitness ^4,5^. Deciphering how these spatial niches shape cellular behavior is critical for elucidating biofilm function, pathogenicity, and resistance ^6,7^.

Resolving transcriptional activity within biofilm architectures remains technically challenging, as methods must preserve native structure while detecting low-abundance, transient transcripts. Despite these challenges, confocal laser scanning microscopy (CLSM) combined with fluorescence *in situ* hybridization (FISH) has advanced the structural and compositional characterization of biofilms by mapping the spatial distribution of phylogenetically distinct populations based on ribosomal RNA (rRNA) ^8,9^. Extending these conventional approaches to mRNA detection is severely hindered by limited DNA probe penetration and RNA secondary structures ^10–12^. Recent image-based approaches have nevertheless advanced bacterial spatial transcriptomics by enabling highly multiplexed *in situ* mapping of endogenous bacterial transcripts while retaining cellular context ^13,14^. These methods, however, typically rely on tiled sets of tens of DNA probes per transcript, multiple rounds of sequential hybridization and imaging, or signal amplification, together with custom fluidics and large probe libraries ^13–15^. Such requirements limit accessibility, and repeated hybridization cycles are difficult to reconcile with fragile, hydrated three-dimensional biofilm structures. Genetically encoded transcriptional reporters offer an alternative, but they require strain engineering, report promoter activity rather than endogenous transcripts, and cannot be applied directly to clinical or environmental isolates ^6^. Unlike DNA probes, peptide nucleic acid (PNA) probes, synthetic nucleic acid analogues with neutral backbones, overcome these barriers, enabling high-affinity, mismatch-sensitive hybridization and enhanced diffusion through complex biofilm structures ^16–18^. While it has been widely used for rRNA-based species identification^19,20^ to our knowledge, PNA-FISH has not been applied to spatially resolve mRNA transcripts within complex three-dimensional matrices. Furthermore, the superior thermodynamic stability of PNA enables robust multiplex detection in a single hybridization step, significantly reducing workload and technical variability compared to sequential singleplex approaches ^19,21^.

Here, we present a multiplex mRNA PNA-FISH workflow for quantitative, *in situ* mapping of targeted transcriptional profiles within intact three-dimensional biofilm architectures. The approach combines short, directly labelled PNA probes designed to hybridize under a common condition, enabling simultaneous detection of a defined transcript panel in a single hybridization step without repeated rounds of hybridization or signal amplification. This demonstrates that a single directly labelled probe per target, without amplification, provides a quantitative *in situ* readout of bacterial mRNA, and it is the first application of PNA-FISH to mRNA within three-dimensional biofilms. We deliberately chose *Legionella pneumophila* because its biphasic life cycle provides a well-characterized transcriptional ground truth, allowing the imaging readout to be tested against established biology across planktonic and biofilm growth conditions. *L. pneumophila* exhibits a tightly regulated biphasic life cycle, transitioning between replicative, non-virulent (exponential phase) and transmissive, virulent (stationary phase) states ^22–25^. This switch is accompanied by the induction of transmission- and virulence-associated genes such as macrophage infectivity potentiator (*mip*) ^26,27^ and the flagellar sigma factor (*fliA*) ^28,29^ (**Supplementary Table 1**).

This well-characterized transcriptional programme therefore provides a biological benchmark for evaluating the quantitative performance of the imaging readout. We show that multiplex mRNA PNA-FISH reproduces the expected transcriptional patterns across planktonic and biofilm populations while preserving relative transcript information between singleplex and multiplex formats. Applied to flow-cell biofilms, the method reveals that sessile populations do not recapitulate the planktonic program and that, in developed biofilms, transcriptional differences track cellular aggregation state and are independent of distance from the fluid interface. Together, these results establish multiplex mRNA PNA-FISH as an accessible, targeted approach for spatially resolved microbial transcriptomics and the systematic, hypothesis-driven investigation of gene-expression patterns across complex microbial communities.

## Results

### Multiplexed mRNA PNA-FISH Workflow for Quantifying Gene Expression

To characterize the spatial dynamics of gene expression within complex microbial architectures, we established a robust, versatile mRNA PNA-FISH workflow. The overall workflow is summarized in **Fig. 1**, comprising: **(1)** *in silico* probe design, **(2)** optimization in planktonic cells and monospecies biofilms, **(3)** image acquisition, **(4)** image analysis, and **(5)** quantitative validation by RT-qPCR.

**Fig. 1.**
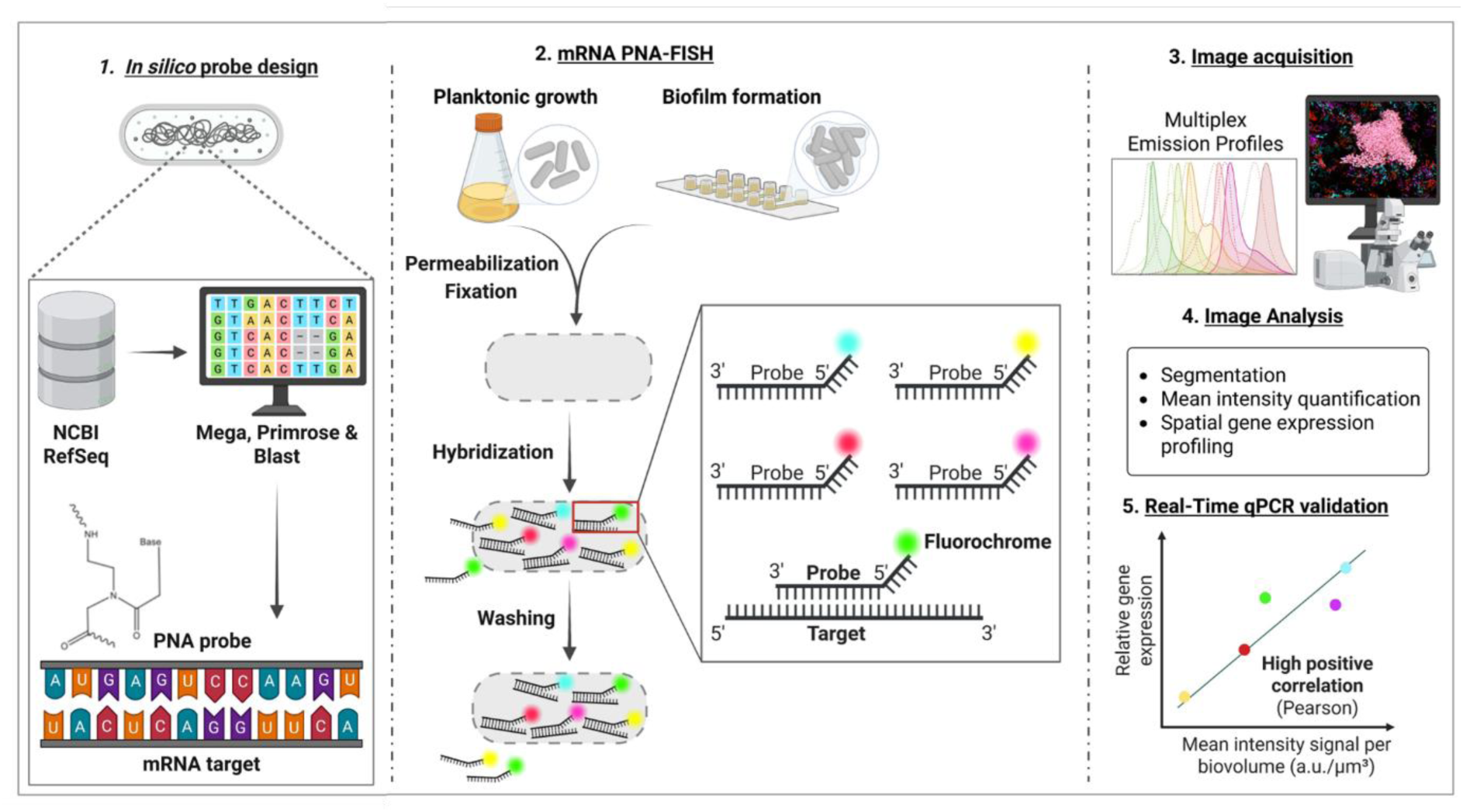
| Workflow for spatial quantification of gene expression in *Legionella pneumophila* biofilms using mRNA PNA-FISH. **(1)** *in silico* design of PNA probes targeting specific mRNA transcripts, **(2)** *in vitro* experimentation for planktonic cells and monospecies biofilms, **(3)** imaging acquisition, **(4)** quantitative analysis, and **(5)** correlation analysis between mRNA PNA-FISH mean intensity and relative gene expression (RT-qPCR). Created with <u>BioRender.com</u>.

The workflow begins with a stringent *in silico* design step, during which candidate target sequences are identified by aligning the transcriptome of interest against comprehensive genomic databases, ensuring high specificity and accuracy.

To enable single-step multiplexing, probes were selected using defined physicochemical criteria: oligomer length (14 -16 nt), GC content (45 - 60%), and Gibbs free energy (*△G*°-13 to -20 kcal mol⁻¹) ^30,31^. Application of these constraints yielded a probe set with closely matched melting temperatures (Tm), a prerequisite for achieving simultaneous hybridization of all targets under uniform conditions ^16^. A common hybridization temperature (60 °C) was established during single-probe optimization and applied throughout all multiplex experiments. Regarding probe chemistry, synthetic PNA oligomers were selected. Due to their uncharged backbone, these probes overcome electrostatic repulsion with the target mRNA, resulting in superior thermodynamic stability compared to conventional DNA analogues ^17,32^.

Importantly, each transcript was targeted by a single probe carrying a single fluorophore (**Supplementary Table 1**). Detection therefore does not depend on tiling tens of probes along a transcript, on signal amplification, or on sequential readout rounds, as required by DNA-based bacterial spatial transcriptomics approaches ^13–15^ This minimal design is enabled by the high affinity and mismatch sensitivity of short PNA probes, and it allows probes to be placed in short, accessible regions of bacterial transcripts, which simplifies panel design and makes each probe an independent, interchangeable module ^16,17^.

Following hybridization, three-dimensional fluorescence images of targeted gene expression were acquired using CLSM. To maximize multiplexing capacity, we implemented a spectral-temporal acquisition strategy that combines simultaneous imaging of spectrally separated fluorophores with sequential acquisition rounds over time, thereby expanding the number of transcripts that can be resolved within a single sample. To resolve fluorophores with overlapping profiles, we integrated sequential scanning with fluorescence lifetime imaging microscopy (FLIM) ^33^. By distinguishing fluorophores by their fluorescence lifetimes in addition to their emission spectra, this strategy prevented signal crosstalk and ensured robust target separation. This dual strategy enabled reliable discrimination even between fluorophores with closely spaced emission bands ^33^. Cross-talk analysis using single-fluorophore-stained controls revealed negligible bleed-through between acquisition channels, confirming spectral specificity (**Supplementary Fig. 1**).

To integrate robust quantitative analysis of biofilm structures into the overall workflow, a segmentation tool embedded in BiofilmQ was used ^34^. This analysis addresses the topological complexity of biofilms by generating a binary biomass mask, which allows fluorescence to be quantified strictly as the mean intensity per unit of biovolume, rigorously decoupling gene expression signals from local variations in bacterial density.

### Singleplex mRNA PNA-FISH in planktonic growth states

To discriminate the physiological transitions driving the *L. pneumophila* biphasic life cycle (**Fig. 2A**), we targeted a defined set of mRNA transcripts. *secE* was selected as an indicator of the replicative state (exponential phase; EP) ^35^, whereas the virulence-associated factors *rpoS* ^36,37^, *mip* ^26,27^, and *fliA* ^28,29^ were used as indicators of the transmissive state (stationary phase; SP) (**Supplementary Table 1** and **Fig. 2A**). Experimental specificity was validated using both a no-probe control and a non-specific probe in all datasets (**Supplementary Figs. 2** and **3**). These controls confirmed the absence of non-specific binding and established a quantitative baseline for intrinsic bacterial autofluorescence. Autofluorescence, which was most pronounced in short-wavelength detection channels (i.e., 405 nm UV-excitation), was systematically subtracted from all datasets to ensure accurate measurement of fluorescence intensity. To capture *in situ* transitions, bacterial cells were grown planktonically in BYE medium for 48 h. Based on the growth profile (**Fig. 2A**), five representative time points were selected (6, 12, 24, 30, and 36 h), capturing the lag, exponential, and stationary phases, providing a robust framework for single-probe validation. Cell viability and growth kinetics were confirmed by CFU quantification, showing standard exponential and stationary phase profiles (**Supplementary Fig. 4**). CLSM analysis revealed distinct temporal fluorescence patterns corresponding to the bacterium’s transition between the replicative and transmissive states, marked by the expression of *secE* and *fliA*/*mip*, respectively (**Fig. 2B**). Quantification of fluorescence intensity normalized to biovolume (a.u./µm³) identified distinct temporal profiles for each targeted transcript (**Fig. 2C**). These patterns of transcript abundance were independently supported by relative expression analysis using RT-qPCR (**Fig. 2D** and **Supplementary Fig. 5**). To validate the accuracy of the imaging-based method, we compared the PNA-FISH mean fluorescence intensity per biovolume with relative transcript abundance measured by RT-qPCR. Individual correlation analyses confirmed this concordance across all probes (16S: R² = 0.8261; *secE*: R² = 0.9358; *rpoS*: R² = 0.8105; *mip*: R² = 0.8666; and *fliA*: R² = 0.8866; **Fig. 2D**). Multiple regression analysis revealed a robust linear correlation across all target genes (R²=0.8891; **Supplementary Fig. 6),** demonstrating that PNA-FISH signal intensity provides a reliable quantitative measure of relative mRNA abundance *in situ*.

**Fig. 2.**
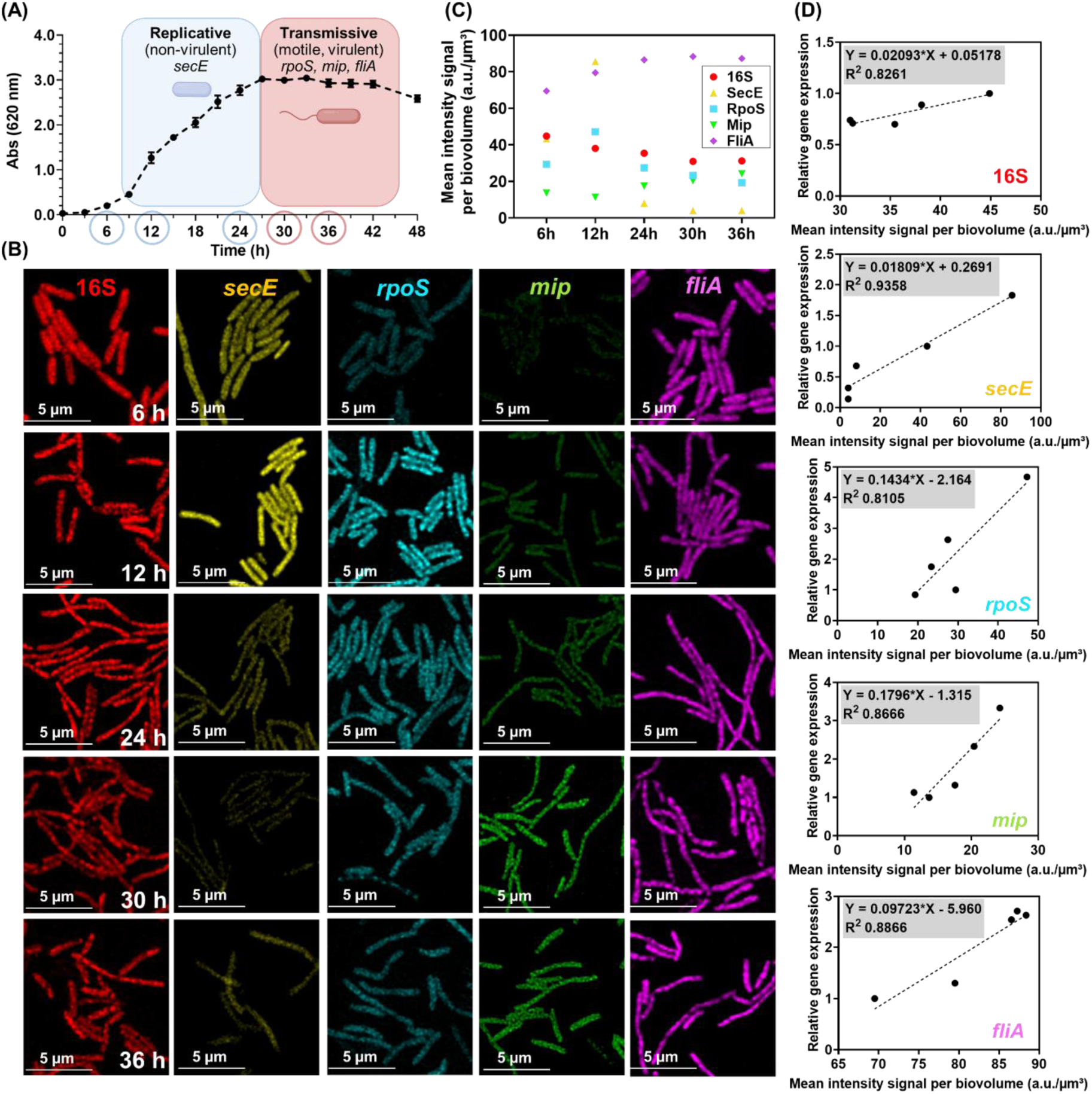
| Quantitative and spatiotemporal analysis of gene expression in *L. pneumophila* by mRNA PNA-FISH. **(A)** *L. pneumophila* planktonic growth on BYE broth over 48 h. The replicative phase (blue box) corresponds to a non-virulent phase and the transmissive phase (red box) to a virulent/motile phase. Error bars represent standard deviation. **(B)** Confocal microscopy analysis of planktonic *L. pneumophila* hybridized at 60 °C at different time points (6, 12, 24, 30, and 36 h) for 16S, *secE*, *rpoS*, *mip*, and *fliA.* **(C)**. Temporal dynamics of target transcripts (16S, *secE*, *rpoS*, *mip*, and *fliA*). Data represent the mean fluorescence intensity per biovolume (a.u./µm³). **(D)** Scatter plot showing strong linear correlation between average mRNA PNA-FISH signal intensity and transcript abundance measured by RT-qPCR. Dashed lines represent the linear regression for each gene. The coefficient of determination (R²) is indicated for each regression.

Several features of these data indicate that the signals originate from the targeted transcripts, excluding non-specific binding or hybridization to genomic DNA as major contributors. First, all targeted genes are single-copy chromosomal genes, so probe binding to genomic DNA would produce broadly similar temporal trends for all targets. Instead, each transcript displayed a distinct profile, and *secE* changed in the opposite direction to *fliA* and *mip* within the same cultures (**Fig. 2**). Second, these gene-specific profiles matched the independently measured RT-qPCR trajectories for every target (**Supplementary Fig. 2D** and **Fig. 5**). Third, they reproduced the expression dynamics expected from the extensively characterized biphasic program of *L. pneumophila* ^22–25^. Fourth, no-probe and non-target probe controls showed no signal above the autofluorescence baseline ( **Supplementary Figs. 2** and **3**). The convergence of an imaging readout, an independent molecular method, and established biology therefore provides orthogonal support for target specificity. The protein secretion gene *secE* exhibited peak expression exclusively during the exponential growth phase at 12 h (∼85.0 a.u./µm³). Matching this profile, RT-qPCR analysis revealed maximal *secE* transcript abundance at the same time point (1.82-fold change), followed by a drastic reduction (fold change < 1.0) in all subsequent stationary phase time points (24 - 36 h). This profile validates *secE* as an indicator for the replicative physiological state, consistent with its role in active growth and protein secretion ^35^.

**Fig. 3.**
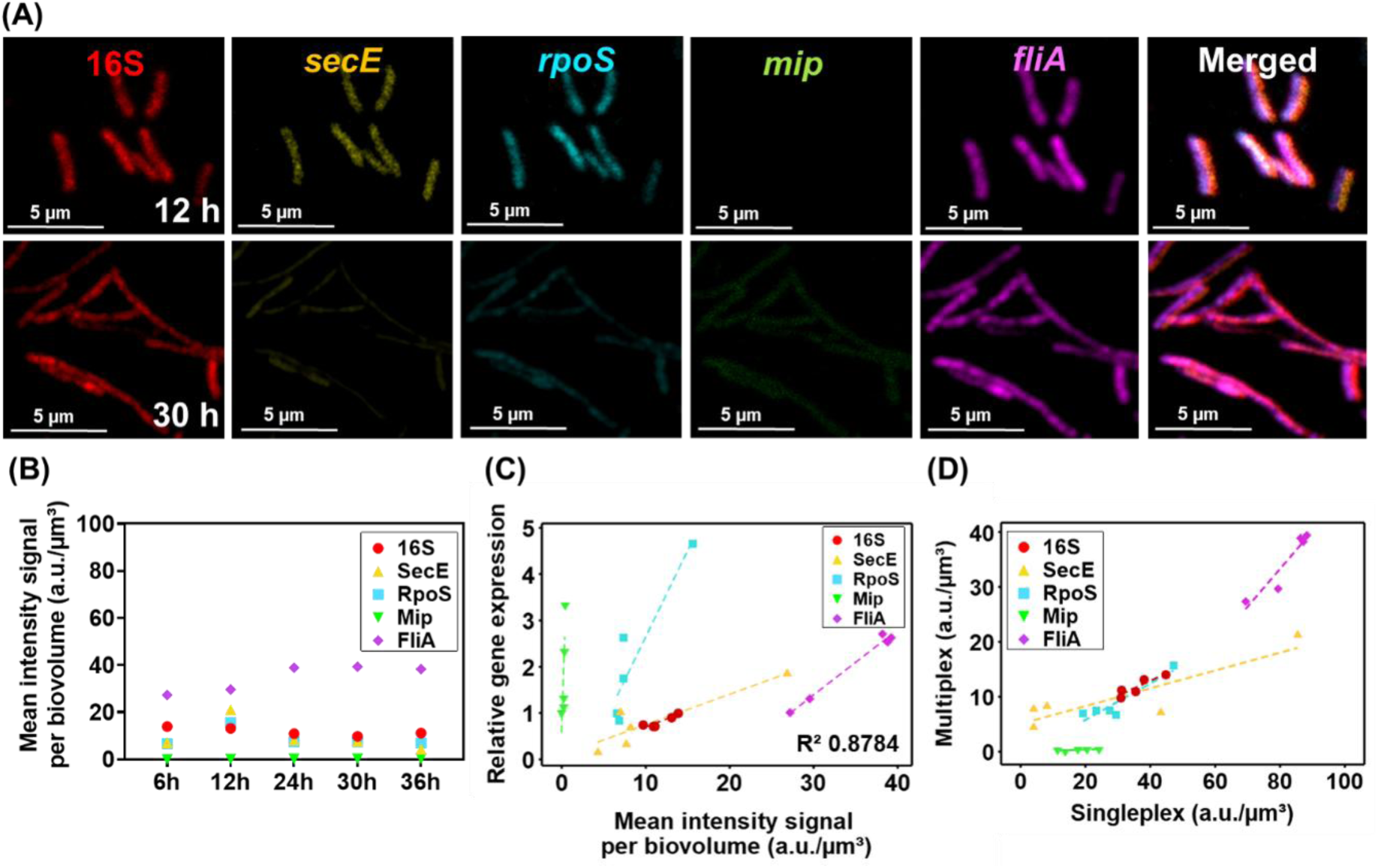
| Multiplex visualization of the transcriptional switch driving the *L. pneumophila* biphasic life cycle. **(A)** Confocal microscopy analysis of planktonic *L. pneumophila* hybridized simultaneously with the mRNA PNA-FISH. **(B)** Quantitative dynamics of gene expression in planktonic cells. Data represent the mean fluorescence intensity per biovolume (a.u./µm³). **(C)** Scatter plot demonstrating a high linear correlation (R² = 0.8784) between the multiplex mRNA PNA-FISH mean signal intensity per biovolume and relative gene expression measured by RT-qPCR. **(D)** Scatter plot comparing Multiplex versus Single-probe signal intensities. The distinct linear proportionality confirms that relative expression trends are preserved in the multiplex format, despite absolute signal attenuation (deviation from the 1:1 diagonal).

Conversely, the transmissive phase was characterized by the strong upregulation of virulence-associated genes. FliA, the flagellar sigma factor, showed a robust induction in the stationary phase, peaking at 30 h (∼88.0 a.u./µm³). This accumulation was reflected by RT-qPCR, which detected a sustained ∼2.7-fold increase in mRNA abundance (**Supplementary Fig. 5**), consistent with the established role of *fliA* in motility ^28,29^. Similarly, the macrophage infectivity potentiator *mip* gradually increased during late growth, reaching its maximal levels at 36 h (∼24.0 a.u./µm³), supported by a significant ∼3.3-fold transcriptional induction (**Supplementary Fig. 5**).

### Multiplex mRNA PNA-FISH for simultaneous visualization of transcript abundance

To evaluate the ability of the method to simultaneously detect multiple mRNA targets, multiplex mRNA PNA-FISH was applied across the planktonic growth kinetics described above. Two time points, 12 and 30 h, were specifically selected as they represent the peak of the exponential and stationary phases, respectively, thereby effectively capturing the biphasic transition (**Fig. 2A**).

CLSM imaging revealed a distinct shift in population-level transcriptional profiles between growth phases (**Fig. 3A**). At 12 h, cells exhibited high fluorescence intensities for the replicative marker *secE* (∼20.0 a.u./µm³) and the stress regulator *rpoS* (∼17.0 a.u./µm³). In contrast, the 30 h population showed a severe decrease in *secE* fluorescence (∼5.0 a.u./µm³), accompanied by a significant induction of the transmissive marker *fliA* (∼40.0 a.u./µm³).

Importantly, most individual channels demonstrated clear, target-specific labeling (**Fig. 3A**). Notably, the spectral narrowing required for multiplex acquisition reduced absolute signal intensities relative to singleplex conditions. As captured in the CLSM images, this optical constraint was particularly noticeable for *mip*, rendering its visual signal weak. The small size of *L. pneumophila* (typically ∼1–2 µm in length), combined with the optical diffraction limit precludes the spatial resolution of individual transcripts within a single cell. We therefore used total fluorescence intensity per biovolume as a measure of relative transcript abundance ^38,39^. Quantitative analysis of target-specific fluorescence intensities per biovolume confirmed these temporal expression patterns (**Fig. 3B**).

To validate the method’s robustness, multiplex mRNA PNA-FISH measurements were compared with relative transcript abundance as described above. A strong correlation was once again observed between multiplex signal intensities and bulk RT-qPCR measurements across targets (R² = 0.8784; **Fig. 3C**). In addition, direct comparison of multiplex and singleplex fluorescence intensities revealed distinct linear proportionality for key transcripts, demonstrating that multiplex detection preserves relative transcriptional information obtained from single-probe assays (**Fig. 3D**). This confirms that the multiplex mRNA PNA-FISH reliably reports relative transcriptional changes, providing biological information equivalent to singleplex assays, with the added advantage of high-throughput, simultaneous detection.

### Temporal dynamics of gene expression in biofilms and planktonic populations

Following the validation of the mRNA PNA-FISH approach within planktonic cultures, which confirmed probe specificity and demonstrated that fluorescence intensities accurately reflect relative gene expression during the biphasic cycle, we extended its application to resolve the spatial organization within the complex three-dimensional architecture of biofilms.

Unlike planktonic populations, which follow a linear transition from replication to transmission, biofilms generate chemical gradients that may support simultaneous and diverse physiological states ^3,40^. To capture the physiological heterogeneity of the attached phase, biofilms were cultivated in a flow cell system under continuous hydrodynamic flow, allowing for monitoring development from initial adhesion (1 h) to biomass accumulation (day two) and full structural maturation (day five). As expected, CLSM showed a progressive quantitative increase in architectural complexity over time (**Fig. 4A**). At 1 h, cells were predominantly organized as a sparsely attached single layer (∼2 µm). By day two, the biomass was largely characterized by a dense, confluent layer, although initial three-dimensional structures up to ∼4 µm in height were already visible. By day five, the biofilm expanded extensively, maturing into multi-layered three-dimensional aggregates with a maximum thickness of ∼14 µm. Importantly, viable cell counts obtained by CFU plating remained consistently high (∼10⁸ CFU/cm²) across all developmental stages (1 h, days two and five), confirming that the visualized structural network represents a continuously viable population (**Supplementary Fig. 7**).

**Fig. 4.**
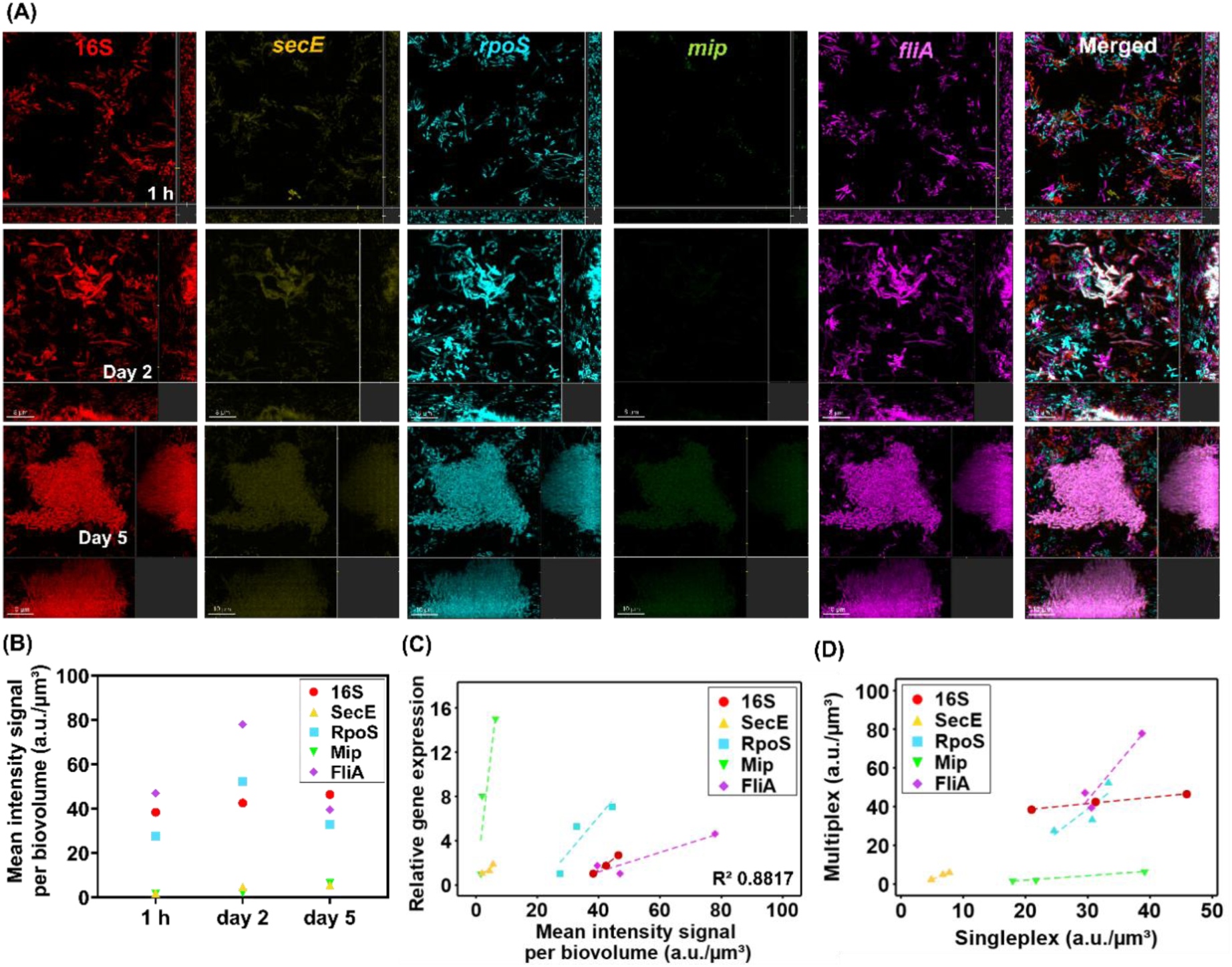
| Spatiotemporal profiling of gene expression during *L. pneumophila* biofilm development. **(A)** Confocal microscopy images for the detection of target transcripts (16S*, secE, rpoS, mip, fliA*) at 60 °C, after 1 h of initial attachment and on days two and five under flow conditions. **(B)** Quantitative dynamics of gene expression in biofilms. Data represent the mean fluorescence intensity per biovolume (a.u./µm³). **(C)** Scatter plot showing a strong multiple correlation (R² = 0.8817) between average mRNA PNA-FISH signal intensity and transcript abundance measured by RT-qPCR for all genes. **(D)** Scatter plot comparing Multiplex versus Singleplex signal intensities. The distinct linear proportionality confirms that relative expression trends are preserved in the multiplex format, despite absolute signal attenuation (deviation from the 1:1 diagonal).

By day two, the developing biofilm layer exhibited a distinct transcriptional signature. Quantitative analysis of the multiplex mRNA PNA-FISH signals (**Fig. 4B**) indicated that the replicative marker *secE* remained at constitutively lower levels of intensity (∼4.0 a.u./µm³). This represents an ∼9-fold reduction compared to the pronounced exponential peak observed in planktonic cultures (**Figs. 2** and **4**). Furthermore, this low transcript abundance within the biofilm was strongly corroborated by RT-qPCR analysis (R² = 0.8817) (**Fig. 4C**). Concurrently, at this colonization stage, the flagellar sigma factor *fliA* (∼70.0 a.u./µm³) and the master stress regulator *rpoS* (∼40.0 a.u./µm³) were induced, reflecting a coordinated activation of motility and stress-response pathways during initial biofilm establishment ^35,41^. As the biofilm developed into distinct clusters on day five, the transcriptional behavior of *secE* exhibited a significant divergence from planktonic populations. Whereas planktonic cells showed a severe decrease in *secE* fluorescence following the exponential phase (<8.0 a.u./µm³), biofilm populations maintained constitutively low levels of *secE* (∼5.0 a.u./µm³) throughout biofilm maturation (1 h, days two and five). This suggests that the attached cells are not driven by a highly replicative subpopulation, a trend supported by RT-qPCR data showing limited *secE* transcript abundance (∼1.7-fold) (**Supplementary Fig. 8A**). Despite the reduced absolute fluorescence inherent to multiplexing described earlier, *in situ* multiplex mRNA PNA-FISH analysis successfully captured the progressive upregulation of the macrophage infectivity potentiator *mip*, throughout biofilm formation. Fluorescence intensity for *mip* reached its maximal level (∼7.0 a.u./µm³) in the developed five-day biofilm. This spatial observation was strongly supported by bulk RT-qPCR data, which identified *mip* upregulation as the most significant transcriptional transition at day five (**Supplementary Fig. 8A**). This *mip* induction coincides with the transition to a virulent, transmissive state, supporting increased *mip* expression as a feature associated with late-stage *Legionella* biofilms ^26^. In contrast, expression levels of the early adaptive transcripts *fliA* and *rpoS* declined on day five, consistent with maturation into a fully sessile lifestyle ^28,42^.

The multiple regression between the multiplex mRNA PNA-FISH quantification and bulk RT-qPCR data in biofilms demonstrated a strong correlation (R² = 0.8817, **Fig. 4C**). In addition, direct comparison of multiplex and singleplex fluorescence intensities from independent flow cell biofilm assays demonstrated linear proportionality for key transcripts, confirming that multiplex detection retains the relative transcriptional relationships observed in singleplex assays (**Fig. 4D**). Individual correlation analyses further confirmed this agreement across all target genes (**Supplementary Fig. 8B**), validating the method’s capability to profile gene expression quantitatively even within optically complex biological matrices such as biofilms. Complementary single-probe imaging further confirmed that the spatial distribution patterns of individual transcripts were preserved in the multiplex format (**Supplementary Fig. 8C**).

### Spatial distribution and heterogeneity of gene expression during biofilm development

To investigate the spatial organization of gene expression during biofilm development, we performed high-resolution vertical profiling of *L. pneumophila* biofilms at early (day two) and developed (day five) stages (**Fig. 5A** and **5B**). To accurately map this vertical architecture, transcript abundance was quantified as a function of distance from the outer biofilm-fluid interface (defined as 0 µm) downwards into the deep, substrate-attached interior. To account for non-uniform cell density throughout the biofilm depth, target mRNA PNA-FISH intensities at each depth interval were normalized to 16S rRNA intensity and subsequently scaled to a relative expression index ranging from 0 to 1.

**Fig. 5.**
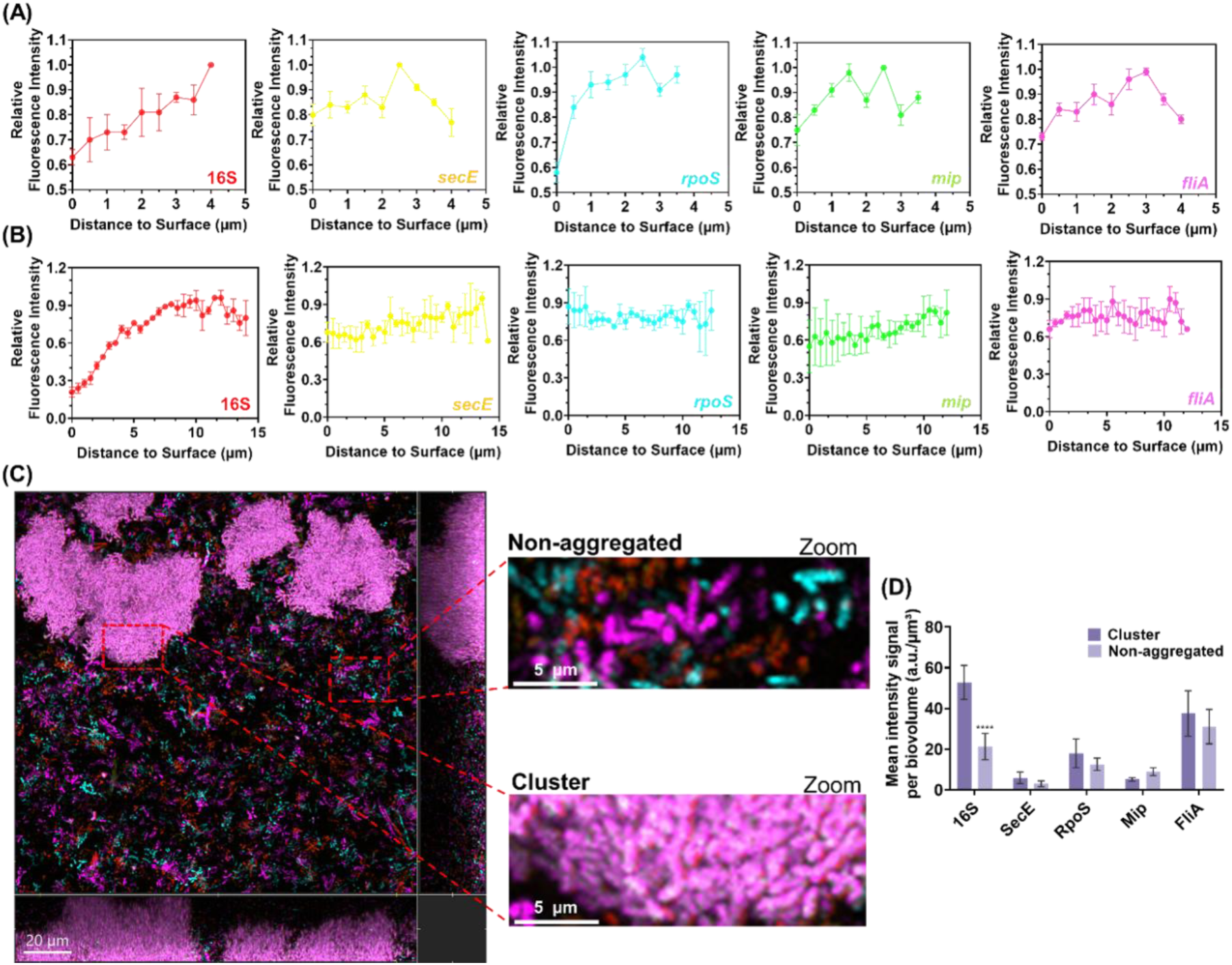
| Spatial organization and phenotypic decoupling during biofilm maturation. Vertical spatial profiling of transcript abundance at day two **(A)** and day five **(B)** for 16S, *secE*, *rpoS*, *mip,* and *fliA*. **(C)** Representative confocal image at day five and zoomed views of non-aggregated and clustered cells. **(D)** Mean fluorescence intensity per biovolume in day five clusters versus dispersed cells (a.u./µm³). Error bars represent standard deviation. Significant differences between clusters and dispersed cells for all the genes (*p* < 0.0001).

At day two (**Fig. 5A**), *L. pneumophila* biofilms exhibit a non-confluent architecture comprising microcolonies up to 4 µm in local thickness. The 16S rRNA profile indicates a higher biomass density at the basal layer. The mean spatial profiles for the targeted transcripts (*secE*, *rpoS*, *mip*, and *fliA*) show expression variations at intermediate depths (1.5 to 3.0 µm). As detailed in the individual microcolony profiles (**Supplementary Fig. 9A**), these spatial patterns exhibit localized transcriptional heterogeneity among discrete microcolonies during initial surface colonization.

By day five (**Fig. 5B**), the biofilm transitions into an established, spatially complex community, with clusters around 15 µm in local thickness (individual cluster profiles are detailed in the **Supplementary Fig. 9B**). The 16S rRNA signal increases continuously with depth, reaching peak intensity in the basal regions of the matrix (10-14 µm), reflecting high basal biomass density. In contrast to the localized heterogeneity observed at day two, the mean spatial expression profiles for *secE*, *rpoS*, and *fliA* remain stable across the biofilm depth. The expression profile for *mip* displays a gradual increase towards the deeper layers (∼1.5-fold), although no localized peaks were observed. The uniform vertical distribution of these functional markers across the analyzed clusters indicates that absolute distance from the fluid interface does not drive transcriptional variation at this biofilm development stage.

To determine how spatial localization influences the physiological state of non-aggregated cells, the mean fluorescence intensity per biovolume was quantified in the day five established clusters compared to the surrounding non-aggregated cells (**Fig. 5C** and **5D**). 16S rRNA intensity was significantly higher in established clusters (∼53.5 a.u./µm³) than in dispersed cells (∼21.0 a.u./µm³, *p* < 0.0001), suggesting a compounding effect of decreased localized biomass and a shift in physiological state in the non-aggregated population. Despite this reduction in signal intensity, the transcription of virulence and motility markers was detected in both subpopulations. Non-aggregated cells exhibited levels of *fliA* (∼31.0 a.u./µm³) with a 1.2-fold difference compared to those in clusters (∼38.0 a.u./µm³). Furthermore, *mip* expression was 1.8-fold higher in the dispersed population (∼9.0 a.u./µm³) relative to clusters (∼5.0 a.u./µm³).

These signals diverged in opposite directions within the same images and under normalization per biovolume, a pattern that cannot be attributed to lower cell density or segmentation effects in dispersed regions, which would reduce all signals proportionally. Together, these data demonstrate that, within the same biofilm, aggregated and non-aggregated subpopulations differ in ribosomal content and in the relative abundance of virulence-associated transcripts, consistent with metabolic compartmentalization while preserving a coordinated virulence transcriptional profile in the dispersed population.

## DISCUSSION

Deciphering how gene expression is spatially organized within the intact three-dimensional architecture of biofilms remains a central challenge in microbial ecology. Bulk transcriptomics captures population-level trends, averaging across coexisting physiological states, but fails to resolve the spatial microenvironments that govern individual cell behaviour ^6^. Here, we established a multiplex mRNA PNA-FISH workflow that directly addresses this barrier by combining short, directly labelled PNA probes designed for a common hybridization condition with a spectral-temporal CLSM acquisition strategy and biovolume-normalized segmentation, enabling simultaneous quantitative spatial profiling of a defined transcript panel within preserved three-dimensional biofilm architecture.

Current bacterial spatial transcriptomics strategies, including par-seqFISH applied to *P. aeruginosa* biofilms, smFISH applied to *V. cholerae* biofilms, bacterial-MERFISH, and RAINBOW-seq, have greatly advanced the spatial mapping of bacterial gene expression. Highly multiplexed approaches have enabled the simultaneous detection of hundreds of transcripts and serve as powerful discovery tools for global transcriptional mapping ^13,14,43,44^. Signal-amplified HCR-FISH has also enabled the simultaneous visualization of multiple target RNAs in *P. aeruginosa* aggregates ^15^. However, these methods typically require extensive probe sets, sequential hybridization rounds, or signal amplification. Most have been applied to thin biofilms, often consisting of a single cell layer, or to static culture systems, or provide limited single-cell or cluster-level resolution in three-dimensional biofilms. In contrast, the neutral backbone chemistry of PNA probes reduces electrostatic repulsion, promoting efficient penetration and hybridization using short probe sequences ^16–19^. In the present workflow, each target is detected using a single directly labelled PNA probe, and all probes are designed to hybridize under a common condition, so a defined transcript panel is detected simultaneously in a single hybridization without signal amplification. Spectral-temporal CLSM provides the multiplex readout, while RT-qPCR provides independent quantitative validation. The method yields an intensity-based measure of relative transcript abundance and is suited to targeted analysis of key regulatory pathways, such as the biphasic life cycle of *L. pneumophila* ^22–25^. Applied to *L. pneumophila* biofilms cultivated under continuous flow, the workflow enables quantitative spatial comparison of targeted transcriptional profiles while preserving three-dimensional biofilm architecture.

Three features distinguish this workflow from existing approaches. First, tiled DNA probe sets require target regions long enough to accommodate many probes, whereas a 14-15 nt PNA probe can be positioned within a short accessible region. This facilitates panel design for short bacterial transcripts and allows new targets to be added without redesigning the panel. Second, because the panel requires neither amplification nor sequential readout, hybridization and washing are performed once, directly within the flow cell, avoiding repeated fluid exchanges that could disrupt biofilm architecture. This allowed the workflow to be applied to biofilms cultivated under continuous flow, complementing previous spatial transcriptomics studies performed in static biofilm models, and enabled depth-resolved quantification of multiple transcripts throughout the biofilm volume. Third, the readout relies on standard confocal instrumentation and open-source image analysis, and requires no genetic modification of the organism, so the approach can be applied to unmodified clinical and environmental isolates and readily transferred between laboratories. These features do not replace transcriptome-scale discovery tools. Rather, they fill a complementary niche, analogous to that of RT-qPCR relative to RNA-seq, enabling rapid, hypothesis-driven quantification of defined gene panels *in situ*, including the spatial validation of candidate genes identified by global methods.

Application of mRNA PNA-FISH to planktonic populations accurately captured the temporal biphasic transition, demonstrating high correlation with RT-qPCR measurements (**Figs. 2** and **3**). However, extending the workflow to sessile communities revealed transcriptional complexity in the spatial regulatory network. Unlike planktonic stationary-phase cells, which downregulate replication-associated transcripts, developed biofilms maintained constitutively low *secE* expression alongside progressive upregulation of *mip* (**Fig. 4**). This indicates that attached communities sustain an active virulence program characterized by marked metabolic and transcriptional adaptability, and do not enter a transcriptionally quiescent state ^35^.

The spatially resolved analysis further elucidated the internal organization of these communities, demonstrating a transition from localized transcriptional heterogeneity to structural synchronization during biofilm maturation. At day two, the high intra-sample variance across vertical profiles suggests that early microcolonies undergo independent, highly localized microenvironmental adaptations before establishing a cohesive architecture (**Fig. 5A**). By day 5, the biofilm underwent a structural reorganization. While 16S rRNA levels, indicative of overall ribosomal content, formed a continuous gradient peaking in the deep interior of the developed biofilm matrix (>10 µm), targeted transcripts *secE*, *rpoS*, and *fliA* remained relatively stable across the entire biofilm depth. Although *mip* expression exhibits a gradual increase towards the basal layer, none of these markers display the pronounced spatial polarization seen during early colonization (**Fig. 5B**). This overall uniform spatial distribution demonstrates that absolute distance from the fluid interface does not dictate drastic transcriptional reprogramming within established clusters. This spatial homogeneity reflects the characteristics of the microfluidic flow-cell system, which continuously supplies nutrients and removes waste, which may limit the formation of steep diffusion-driven gradients typical of larger-scale biofilm models ^45^.

Vertical depth alone does not fully account for the transcriptional distribution in developed biofilms; the synchronization of the transmissive switch within these aggregates may be coordinated by local architecture-dependent regulatory networks. Systems such as the Lqs quorum-sensing network and architectural regulators like LvbR are essential for the formation of compact bacterial aggregates and the synchronization of virulence expression ^46,47^. These observations align with the phenotypic heterogeneity described by Personnic et al. (2021), who identified subpopulations of non-growing but metabolically active sessile cells that maintain high virulence expression ^48^. Rather than responding to macro-scale physical gradients, microenvironmental cues retained within the dense EPS matrix are likely integrated through master regulatory networks, such as the LetA/LetS cascade, to govern the transmissive phenotype ^40,49^. Future application of mRNA PNA-FISH targeting components of the Lqs and LetA/LetS systems would directly test this hypothesis and resolve the regulatory architecture underlying the observed spatial transcriptional patterns.

The mean intensity per biovolume quantification reveals that cellular aggregation state, rather than vertical depth, is the main axis of transcriptional differentiation in developed biofilms. A distinct spatial decoupling between metabolic state and virulence programming occurs between aggregated and non-aggregated subpopulations; non-aggregated cells exhibit a marked reduction in global ribosomal content yet maintain motility (*fliA*) and significantly upregulate the macrophage infectivity potentiator (*mip*) relative to dense clusters (**Fig. 5C** and **5D**). The opposing ribosomal content and *mip* signals within the same images rule out density- or segmentation-related artifacts. Moreover, vertical depth, the most obvious alternative driver, did not account for transcriptional differences in developed biofilms (**Fig. 5B**), pointing to aggregation state as an important organizing variable. These quantitative observations indicate that the non-aggregated subpopulation is unlikely to reflect simple nutrient-induced senescence. Instead, it is consistent with a regulated, non-replicative transmissive state resembling the classical stationary phase, but linked to architectural context, not to bulk nutrient depletion. Collectively, these observations suggest that biofilms may continuously generate a dispersed subpopulation that shows reduced ribosomal signal yet transcriptionally primed for host interaction. Non-aggregated cells are those most likely to be released from biofilms into water systems, making this spatial differentiation directly relevant to transmission. This hypothesis, with important potential implications for the transmission dynamics of Legionnaires’ disease, could not be addressed by bulk transcriptomics or by single-reporter approaches, and becomes directly testable *in situ* with the workflow presented here.

The scope of the method should be considered alongside its advantages. As noted above, the workflow measures relative transcript abundance per biovolume, not absolute transcript counts per cell. The current five-target panel is limited by the optical readout. The probe chemistry itself imposes no such constraint. All probes hybridize simultaneously under a common condition, and additional probes can be added without changing the protocol. The present acquisition already combines spectral separation with fluorescence lifetime discrimination. Extending this to full spectral unmixing across the available excitation range, with lifetime-resolved fluorophore pairs within each spectral window, is expected to allow in the order of 8-10 transcripts to be resolved in a single hybridization on commercial confocal systems, without sequential rounds of amplification ^33,50^. Within these bounds, the method delivers what hypothesis-driven studies most often require: quantitative, simultaneous measurement of a defined set of endogenous transcripts, in unmodified cells, within intact three-dimensional structure. As each transcript is labelled with a defined fluorescent signature, future adaptations of the method could extend this barcoding to fluorescence-activated cell sorting (FACS), allowing cells to be separated according to their transcriptional profile for downstream *ex situ* molecular analysis, such as RNA sequencing.

As a benchmark study, we present a multiplex mRNA PNA-FISH workflow for simultaneous spatial mapping of a defined transcript panel within intact three-dimensional biofilm architectures. Relative transcript abundance is consistently preserved across conditions, as supported by strong correlations with RT-qPCR, ensuring robust quantification at individual clusters and subpopulations using standard CLSM instrumentation. Moreover, the underlying design and imaging principles are compatible with high-dimensional fluorescence strategies, including spectral unmixing and fluorescence lifetime-based discrimination ^50,51^. Although demonstrated here using the biphasic transcriptional program of *L. pneumophila*, the workflow relies on a single-step multiplex hybridization strategy and readily customizable PNA probe panels, enabling rapid adaptation to diverse microorganisms, regulatory pathways, and environmental conditions without extensive protocol redesign. By coupling rapid and robust *in situ* transcript detection with preserved three-dimensional architecture, this approach enables routine investigation of spatial gene-expression programs across microbial communities, transforming spatial context from an inferred property into a directly measurable biological variable.

## Methods

### *In silico* PNA probe design for the specific detection of *Legionella pneumophila* genes

For the design of specific probes to target the mRNA of different *Legionella* genes, the National Center for Biotechnology Information (NCBI) RefSeq database was used to access available sequences (http://www.ncbi.nlm.nih.gov). The MEGA (Molecular Evolutionary Genetics Analysis, version 12.1) with the ClustalW algorithm tool was used to align the sequences in the database ^52^. After that, the Primrose software was used to predict all the possible probes ^53^. This prediction was made based on specific criteria, such as a length of 14-16 nt, Tm above 50 °C, a GC content between 45-60%, and Gibbs free energy (*△G*°-13 to -20 kcal mol⁻¹) ^54^. To ensure sequence specificity, candidate probe sequences were subjected to a two-step *in silico* stringency screen. First, sequences were aligned against the complete *L. pneumophila* genome using BLASTn to identify and exclude potential off-target binding sites within the host transcriptome. Second, to evaluate transcriptional orthogonality for potential applications in polymicrobial matrices, sequences were screened against the genomes of phylogenetically distinct species commonly co-inhabiting aquatic environments, including *Pseudomonas aeruginosa* and *Klebsiella pneumoniae*.

The selected sequences were synthesized (Eurogentec, Belgium) with the N-terminus labeled with a fluorophore molecule via a double 8-amino-3,6-dioxaoctanoic acid (AEEA) linker. In addition, a non-specific control probe (targeting *K. pneumoniae*) based on Azevedo et al., 2022 ^50^ was used to validate probe specificity and assess non-target binding at the cellular level. All probe sequences are listed in **Supplementary Table 1.** Experimental specificity and background signal baselines were validated using both a non-specific control probe and control without probe (**Supplementary Figs. 2** and **3**). Intrinsic bacterial autofluorescence, which was predominant in the short-wavelength detection channels, was systematically quantified using the controls without probe and subtracted from all datasets to ensure accurate measurement of fluorescence intensity.

### Bacteria and growth conditions

*L. pneumophila* serogroup 1, ATCC™ 33152, was obtained from the American Type Culture Collection (ATCC; Manassas, VA, USA). The bacteria were grown on filter-sterilized buffered yeast extract (10 g L⁻¹ yeast extract; VWR International, Radnor, PA, USA) (BYE) buffered with 5 g L⁻¹ ACES (N-2-acetamido-2-aminoethanesulfonic acid), 1 g L⁻¹ α-ketoglutarate, monopotassium salt, 2.8 g L^-1^ potassium hydroxide) supplemented with L-cysteine (0.4 g L^-1^) and ferric pyrophosphate (0.25 g L^-1^) (all supplements and buffers were obtained from Sigma-Aldrich, St. Louis, MO, USA) in a secondary container at 37 °C under agitation in an orbital shaker at 110 rpm (Thermo Scientific™ Digital Mini Rotators, Thermo Fisher Scientific, Waltham, MA, USA). The Buffered Charcoal Yeast Extract (BCYE) agar was used to grow the bacteria in a solid medium containing activated charcoal (2.00 g L^-1^) (VWR International, Radnor, PA, USA). The plate cultures were incubated at 37 °C for two or four days. The strain was stored at -80 °C in cryovials containing 2% peptone enriched with glycerol (20% v/v).

### Planktonic kinetic assays

To mimic the biphasic life cycle, an overnight culture was diluted 1:100 in fresh BYE broth to an optical density (OD_620_) of ∼0.03 (∼1*10^8^ cells mL⁻¹) and incubated at 37 °C for 48 h. Growth kinetics were monitored every 3 h using a spectrophotometer (Genesys 20, Thermo Scientific).

### Colony Forming Unit (CFU) Quantification

The number of cultivable cells at 6, 12, 24, 30, and 36 h of planktonic growth was determined by CFUs enumeration. For this, 1 mL of suspension was recovered, and the pellets were washed with distilled water. Serial dilutions were performed, plated onto BCYE, and then incubated at 37 °C for two or four days. The total colony-forming units per mL (log_10_ CFUs mL ⁻¹) were counted after a minimum of 48 h of incubation. For biofilm experiments, the cells were recovered from the channels, and CFUs were quantified according to the same procedure used for planktonic growth. The results were expressed as log_10_ CFU/cm^3^.

### Flow cell biofilm conditions

*L. pneumophila* biofilms were cultivated in Ibidi µ-Slide VI 0.4 flow channels (channel height: 0.4 mm, growth area: 0.6 cm²) (Ibidi, Germany). The inoculum was prepared from an overnight culture diluted in 1:10 BYE medium to a standardized optical density (OD_620_) of ∼0.2 (∼1*10^9^cells mL⁻¹). The cell suspension (150 µL) was inoculated into the channels and incubated under static conditions for 1 h at room temperature (25 °C) to promote initial adhesion. Subsequently, the system was connected to a syringe pump (NE-1600; New Era Pump Systems, Inc., USA) to initiate a continuous laminar flow of fresh medium at a constant rate of 20 µL min⁻¹. This flow rate generated a calculated wall shear stress of τ ≈ 3.3 × 10⁻³ Pa, simulating low-shear environmental niches ^28^. Biofilms were cultivated at room temperature (25 °C) for up to five days, a condition selected to replicate environmental niches, such as engineered water systems and cooling towers, where *L. pneumophila* is known to persist and form biofilms ^55^. To preserve the *in situ* transcriptional profile, fluid flow was terminated, and flow cells were immediately placed on ice for further analysis.

### Probe optimization in planktonic cells

Probe optimization was performed as previously described by the group with some modifications ^20,56^. The bacterial growth procedure was conducted as described previously. The cell suspension was collected at 6, 12, 24, 30, and 36 h, then centrifuged at 6000 × g, 4 °C for 5 min, and immediately placed on ice. After that, the pellets were washed in distilled water, and then the cells were fixed with 500 μL ice-cold 4% paraformaldehyde (PFA) ^13^ and were incubated on ice for 1 h in the dark, and then 500 μL of 50% (vol/vol) ethanol was added and incubated for at least 30 min at -20 °C. After that, 100 μL of cells were centrifuged under the same conditions. Then 100 μL of hybridization solution containing 10% (wt/vol) dextran sulfate, 10 mM NaCl, 30% (vol/vol) formamide, 0.1% (wt/vol) sodium pyrophosphate, 0.2% (wt/vol) polyvinylpyrrolidone, 0.2% (wt/vol) Ficoll, 5 mM disodium EDTA, 0.1% (vol/vol) Triton X-100, 50 mM Tris-HCl (pH 7.5) (all from Sigma-Aldrich, USA) and 200 nM of each PNA probe (Eurogentec, Belgium). The samples were protected from light and incubated for 60 min at different temperatures under study. Following hybridization, the samples were centrifuged under the same conditions, and 500 μL of washing solution, containing 5 mM Tris base, 15 mM NaCl, and 1% (vol/ vol) Triton X (pH 10) (all from Sigma-Aldrich, USA), was added. Finally, 20 μL of cells were dispensed in 5 mm well slides and allowed to air dry, mounted with a drop of nonfluorescent immersion oil (Citifluor™ AF1, Electron Microscopy Sciences, USA), and covered with coverslips. Slides were stored in the dark for a maximum of 24 h before microscopy visualization. Single probes were initially optimized to define hybridization conditions yielding robust signal-to-noise performance across bacterial growth stages. Multiplex hybridization was subsequently performed using the temperature established during single-probe optimization. Additionally, a probe developed by our group, labeled with Alexa Fluor 594 for the detection of *L. pneumophila* rRNA, was used in all experiments ^57^.

### mRNA PNA-FISH in biofilms

After biofilm growth was completed, the FISH protocol described above was applied with some modifications. Cells were fixed directly in each channel with 150 μL of ice-cold 4% PFA and incubated on ice for 15 min. Then, 150 μL of 50% (vol/vol) ethanol was added and incubated for another 15 min at -20 °C. Hybridization was performed by adding 100 μL of hybridization solution containing 200 nM of the PNA probe to each channel, followed by incubation at 60 °C (previously determined as the optimal hybridization temperature) for 60 min, protected from light. After hybridization, 150 μL of washing solution was added, and incubation continued under the same conditions for 30 min. The channels were then rinsed with distilled water to remove any remaining washing solution. Finally, the flow cells were directly observed under the microscope.

### Microscopy analysis

Fluorescence imaging was performed on a STELLARIS 8 Confocal Laser Scanning Microscope (Leica Microsystems, Wetzlar, Germany), equipped with a White Light Laser (WLL) and high-sensitivity Power HyD detectors. Images were captured using an HC PL APO CS2 63x/1.40 oil immersion objective at a resolution of 1024 × 1024 pixels, with a pinhole set to 1.0 Airy unit. To ensure spectral fidelity and prevent signal crosstalk within the multiplex assay, a sequential acquisition protocol was employed. The excitation sequence was ordered as follows: Alexa Fluor 488, ATTO 633, ATTO 550, ATTO 594, and ATTO 405 (**Supplementary Fig. 10**). For fluorophores exhibiting overlapping emission spectra, FLIM was employed to resolve signals based on photon arrival times. The specificity of this spectral separation was rigorously validated using single-probe control samples, which were imaged under acquisition settings identical to those used for the multiplexed samples (**Supplementary Fig. 1**). For each channel, signals were detected exclusively within their designated detection windows, with negligible bleed-through into adjacent spectral channels.

### Image processing and quantitative analysis

Confocal image stacks were processed using LAS X software (version 4.8.1.29271; Leica Microsystems, Wetzlar, Germany). For quantitative analysis, the MATLAB-based tool BiofilmQ was used to perform segmentation using the Otsu thresholding method, and target fluorescence was quantified as the mean intensity per unit of biovolume (a.u./µm³) ^34^. For each experimental condition, a minimum of five independent fields of view (z-stacks) were analysed. All analyses were performed under identical acquisition and thresholding settings to ensure consistency. To assess the vertical stratification of gene expression within clusters, spatial mapping was performed using the *Distance to Surface* metric. In this algorithm, the biofilm surface is defined by a convex hull enclosing the highest Z-centroid values of the segmented biomass, which physically corresponds to the outer biofilm-fluid interface. Consequently, a spatial distance of 0 µm represents cells at the absolute periphery exposed to the bulk liquid, while increasing distance values represent deeper internal layers descending towards the flow cell substratum.

### Analysis of gene expression of *L. pneumophila*

To analyse the gene expression levels of *L. pneumophila* in planktonic and biofilm lifestyles, Quantitative Real-Time PCR (RT-qPCR) studies were performed. Suspensions were prepared, as described previously, and aliquots for planktonic growth were recovered at 6, 12, 24, 30, and 36 h, and for biofilm at 1 h after attachment, two and five days. All experiments were performed using three independent biological replicates. For each biological replicate, two independent aliquots per time point were processed in parallel for RNA extraction and cDNA synthesis. The RNA was extracted using the TRIzol™ Max™ Bacterial RNA Isolation Kit (Thermo Fisher Scientific, USA) according to the manufacturer’s instructions. To prevent potential DNA contamination, samples were treated with RNase-free DNase I (1 U/μL) (Thermo Fisher Scientific, USA). RNA concentration and purity (A_260/280_ ratio ≥ 1.8) were assessed by optical density measurement using the NanoDrop Spectrophotometer (ND 1000, Version 3.8.1, Thermo Fisher Scientific, USA).

For cDNA synthesis, 100 ng total RNA from each independent aliquot was reverse transcribed using the iScript™ cDNA Synthesis Kit (Bio-Rad, USA) following the manufacturer’s guidelines. Quantitative real-time PCR (qPCR) was performed in 96-well microtiter plates using an Applied Biosystems™ QuantStudio™ 7 Real-Time PCR System (Thermo Fisher Scientific, USA) and SsoAdvanced Universal SYBR Green Supermix (Bio-Rad, USA). Each 10 µL reaction contained 2 µL of the normalized cDNA template (30 ng/µL) and 200 nM of each primer.

To ensure accurate relative quantification, standard curves were generated for each primer set using 10-fold serial dilutions to determine amplification efficiency (E) (90% < E < 110%). Specificity was confirmed via melt curve analysis. Strict controls were maintained for each run, including no-reverse transcriptase controls (NRTs) to verify the absence of residual genomic DNA, and no-template controls (NTCs) to monitor reagent contamination. All qPCR reactions were performed in technical triplicate for each of the two cDNA aliquots per biological replicate, and the average relative expression values for each gene were calculated. The primers were designed using the Primer 3 web-based software (http://www.bioinformatics.nl/cgi-bin/primer3plus/primer3plus.cgi) and are listed in **Table 1**. Transcript levels were normalized with 16S rRNA, and the fold change was calculated using the *Pfaffl* method ^58^ (*ratio* = (*E∗target*) *ΔCttarget*(*control*−*sample*) (*E∗ref*) *ΔCtref*(*control*−*sample*.

**Table 1.**
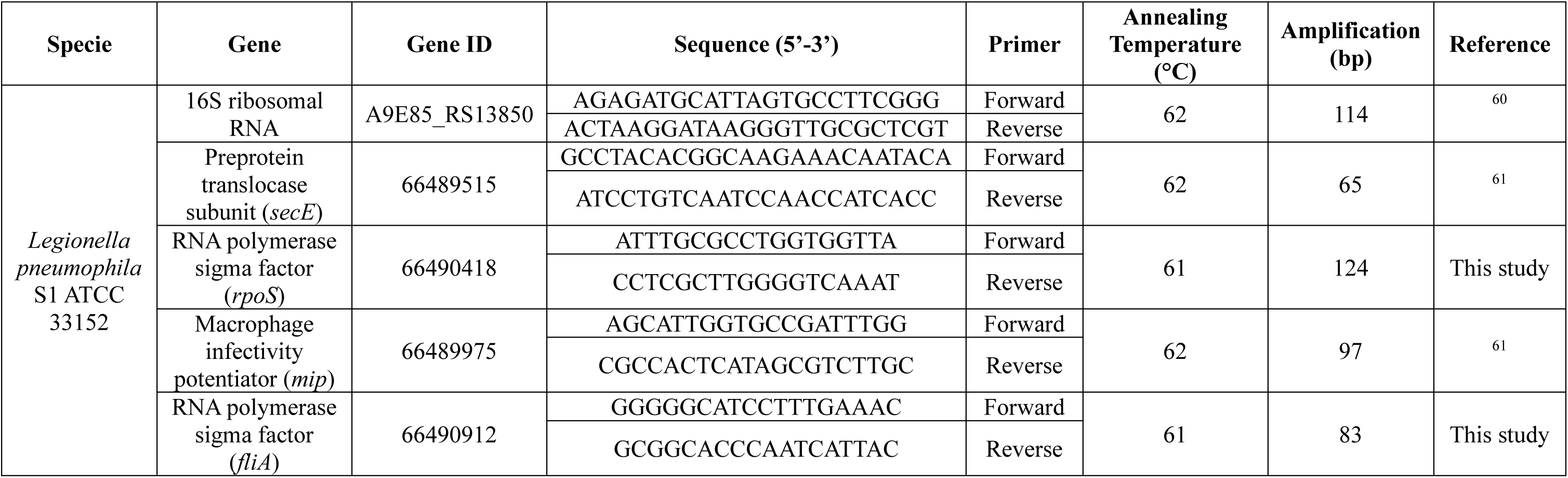
-Primers used for RT-qPCR, with the respective annealing temperature and amplification product.

### Statistical analysis

Experiments were performed in a minimum of three independent biological replicates on independent days. Data are expressed as the mean ± standard deviation (SD). GraphPad Prism 8.4.3® (GraphPad Software, San Diego, CA, USA) was used to perform the variance analyses. Specifically, one-way ordinary ANOVA followed by Tukey’s multiple comparisons test was used to evaluate differences across single experimental variables (CFU counts). For comparisons involving multiple variables (RT-qPCR and mean intensity per biovolume for non-aggregated cells and clusters), a two-way ANOVA was performed, also followed by Tukey’s multiple comparisons test. All statistical tests were performed with a 95% confidence level (*p* < 0.05). The association between mRNA PNA-FISH (mean signal intensity per biovolume (a.u./µm³) and RT-qPCR data was performed using multiple regression using the software R ^59^ with gene as a factor, including the 2-way interaction with signal intensity.

## Author Contributions

Conceptualization, N.F.A., D.M.G. and L.C. Data curation, A.B. and H.J.S. Formal analysis, A.B., H.J.S. and A.P. Funding acquisition, N.F.A., D.M.G. and L.C. Investigation, A.B. Methodology, A.B., H.J.S., A.P., N.F.A., D.M.G. and L.C. Supervision, N.F.A., D.M.G. and L.C. Writing - original draft, A.B. and L.C. Writing - editing, A.B., H.J.S., A.P., C.J.J., N.F.A., D.M.G. and L.C.

## Data Availability

The data supporting the findings of this study are available within the Article and its Supplementary Information. Additional data are available from the corresponding author upon reasonable request.

