## Supplementary Information for "Single-probe, amplification-free multiplex mRNA PNA-FISH maps bacterial gene expression in intact three-dimensional biofilms"

### Supplementary Material

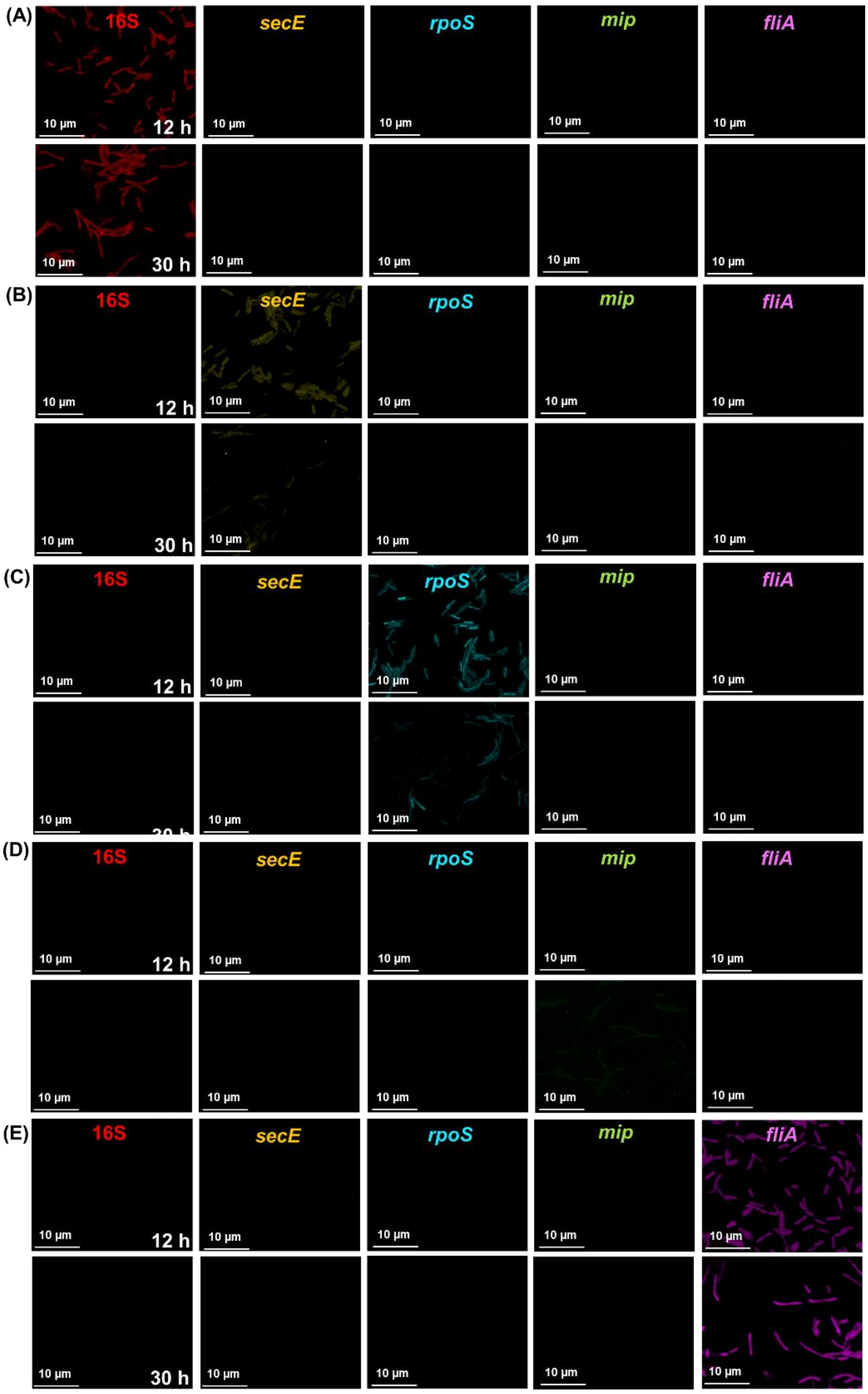


**Supplementary Figure 1 | Single-fluorophore controls validate spectral specificity of multiplex mRNA PNA-FISH**. Confocal laser scanning microscopy (CLSM) images of *L. pneumophila* planktonic cultures at 12 and 30 h and hybridized with singleplex mRNA PNA-FISH targeting **(A)**16S (Alexa Fluor 594), **(B)** *secE* (ATTO 405), **(C)** *rpoS* (ATTO 550), **(D)** *mip* (Alexa Fluor 488), and **(E)** *fliA* (ATTO 633). For each single-probe control, images were acquired across all five detection channels using the same settings applied in multiplex mRNA PNA-FISH experiments.

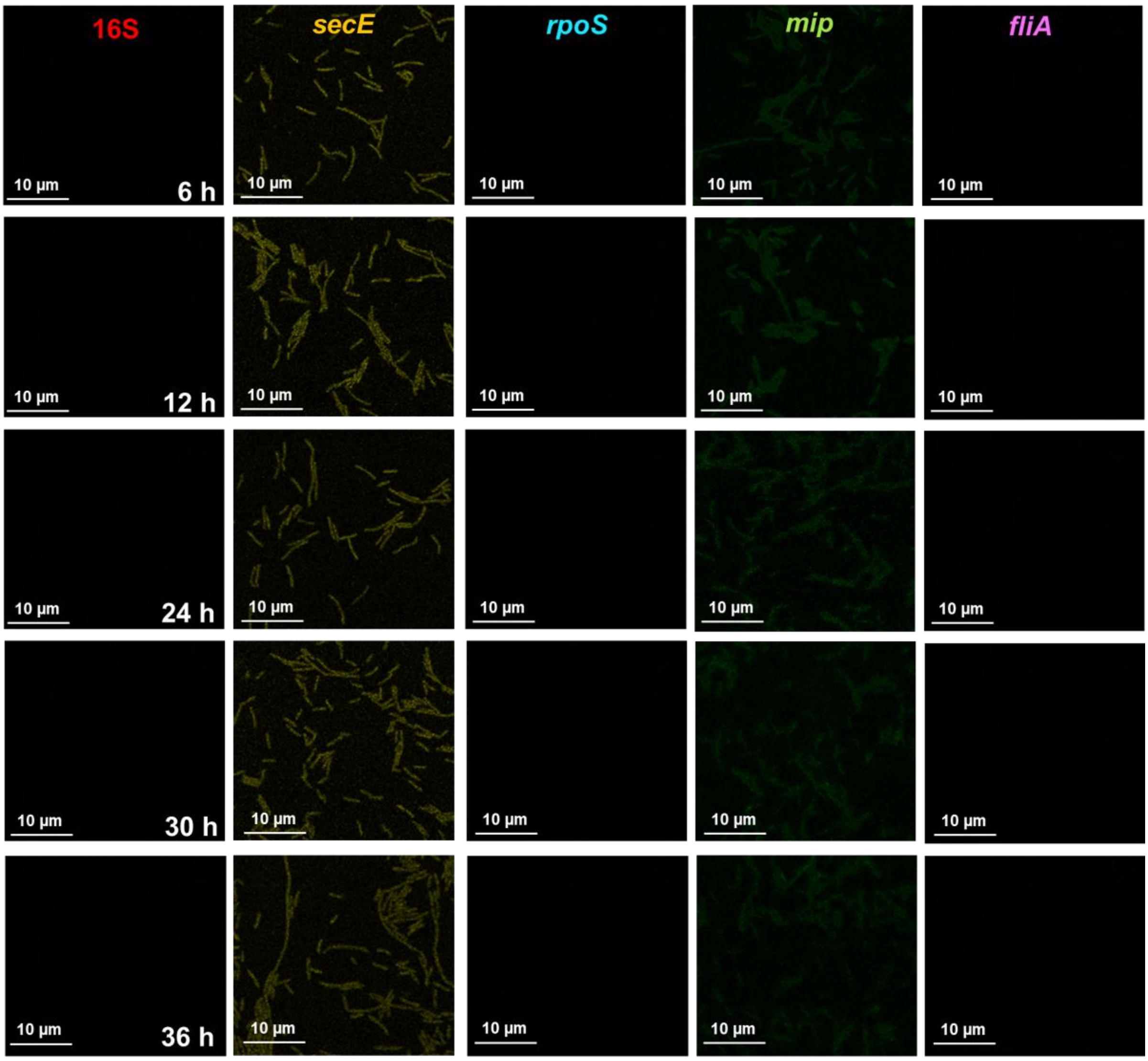


**Supplementary Figure 2 | Background signal controls and autofluorescence profiling of *Legionella pneumophila*.** Confocal laser scanning microscopy (CLSM) images of *L. pneumophila* planktonic growth at different time points (6, 12, 24, 30, and 36 h) without probe. Image acquisition was performed using the same settings as those applied in the specific mRNA PNA-FISH experiments (Alexa Fluor 594 for 16S, ATTO 405 for *secE*, ATTO 550 for *rpoS*, Alexa Fluor 488 for *mip,* and ATTO 633 for *fliA*).

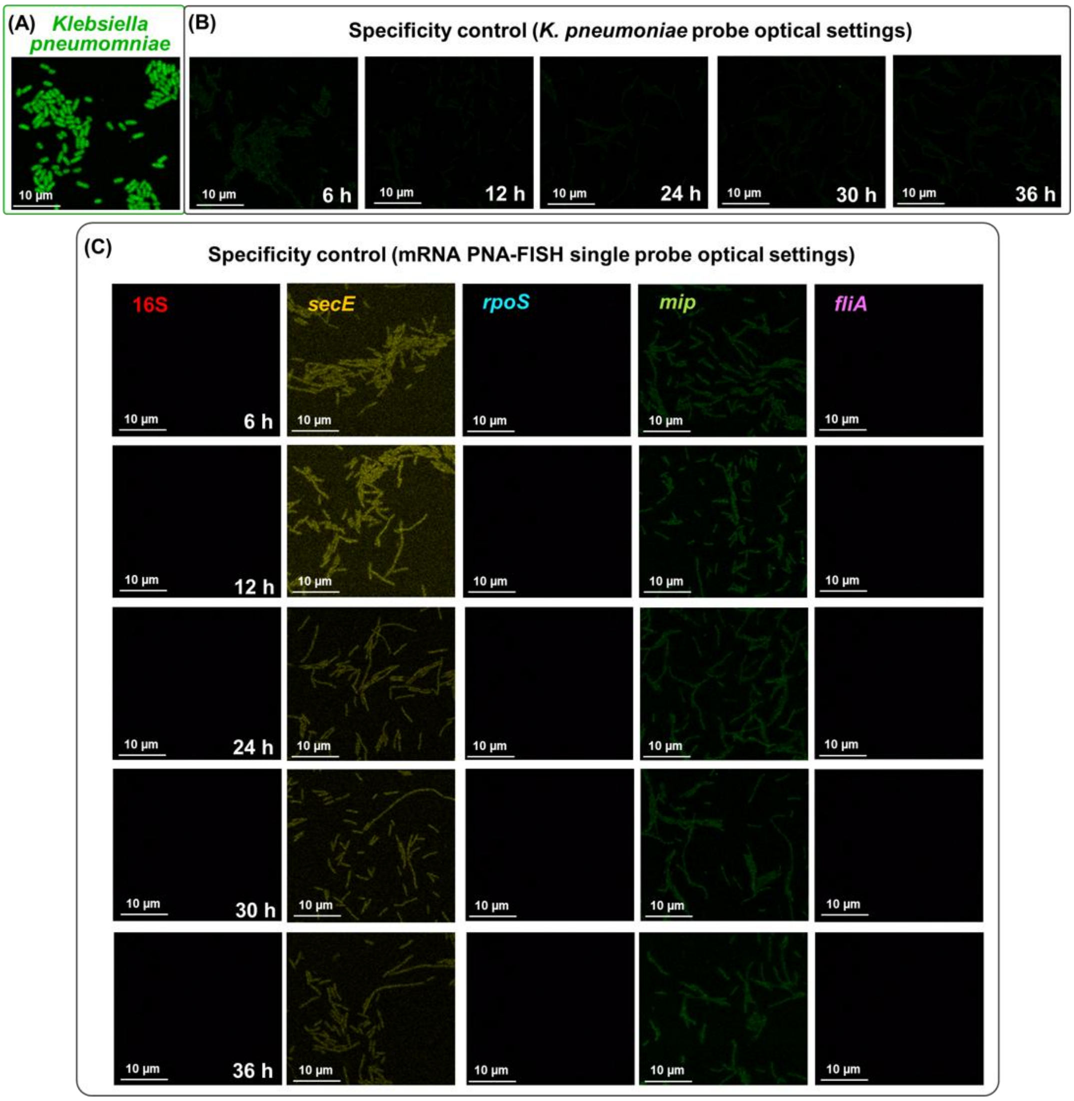


**Supplementary Figure 3 | *Legionella pneumophila* non-specific validation. (A)** Positive control showing the successful hybridization of the non-target *K. pneumoniae* probe to its target species. **(B)** Specificity control demonstrates the absence of cross-hybridization when the non-target probe is applied to *L. pneumophila* across different time points (6, 12, 24, 30, and 36 h), imaged under the non-specific probe's optical settings. **(C)** Specificity control for *L. pneumophila* samples hybridized with the non-target probe imaged using the mRNA PNA-FISH single probe acquisition settings.

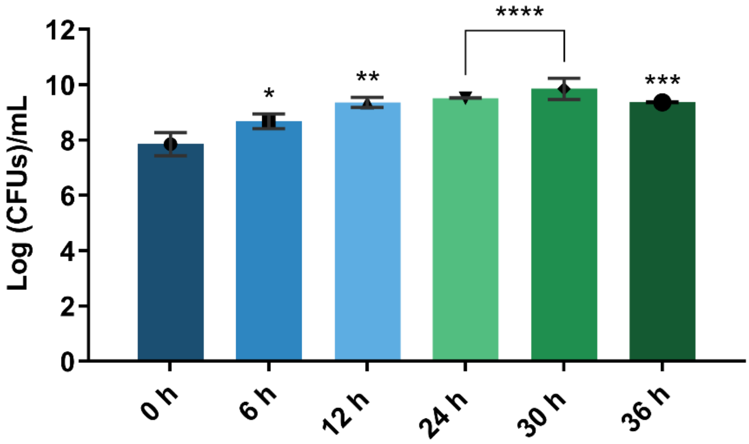


**Supplementary Figure 4 | *Legionella pneumophila* planktonic growth on BYE**. Number of cultivated cells ($\log_{10}$ CFU/mL) at 0, 6, 12, 24, 30, and 36 h. Error bars represent standard deviation. * Significant difference between 0 h and the other timepoints (**p*-value < 0.05, ***p*-value < 0.01, ****p*-value < 0.001, and *****p*-value < 0.0001).

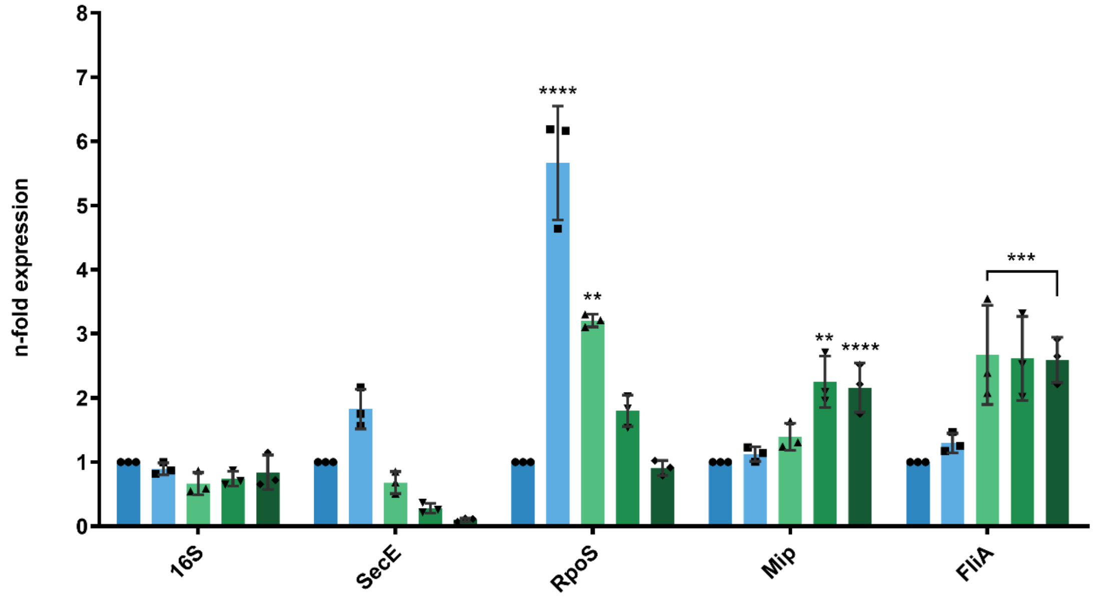


**Supplementary Figure 5 | *Legionella pneumophila* gene expression during planktonic growth on BYE.** Levels of gene expression were quantified using the *Pfaffl* method for *secE*, *rpoS*, *mip*, and *fliA* at 6, 12, 24, 30, and 36 h. Error bars represent standard deviation. * Significant difference between 6 h and the other time points for each gene (***p*-value < 0.01, ****p*-value < 0.001, and *****p*-value < 0.0001).

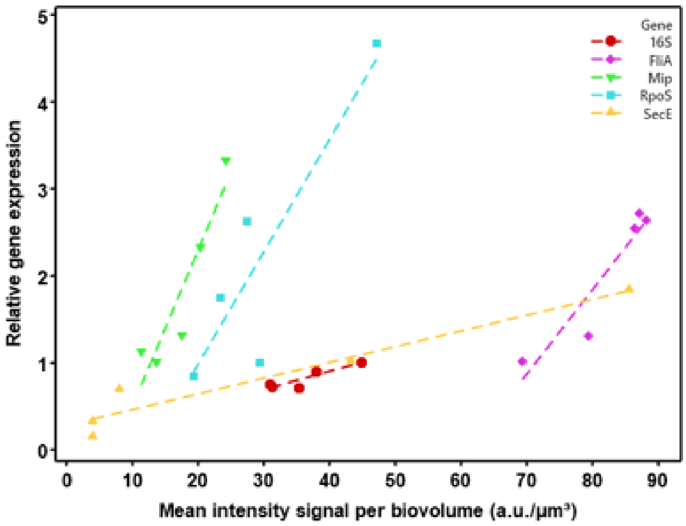


**Supplementary Figure 6 | Global correlation between singleplex mRNA PNA-FISH fluorescence and gene expression during planktonic growth**. Scatter plot illustrates the correlation between mean PNA-FISH fluorescence intensity (normalized to biovolume) and relative transcript abundance measured by RT-qPCR. Data points represent pooled measurements across five targets (16S, *secE*, *rpoS*, *mip*, and *fliA*) at 6, 12, 24, 30, and 36 h. The strong global linear association ($R^2 = 0.8891$) demonstrates a high degree of concordance between the mRNA PNA-FISH signal intensity and RT-qPCR.

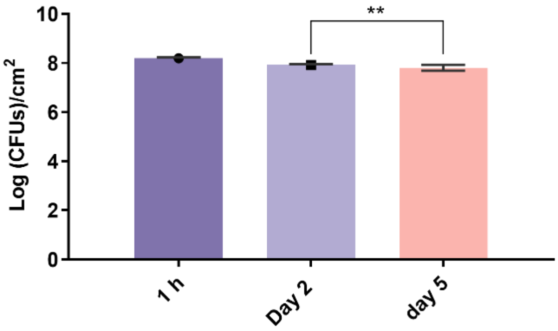


**Supplementary Figure 7 | *Legionella pneumophila* biofilms under flow conditions.** Number of cultivated cells [log10 (CFUs)/cm$^3$] measured at 1 h after initial attachment, and on days two and five. Error bars represent standard deviation. * Significant difference between 1 h and the other time points ($p$-value < 0.01).

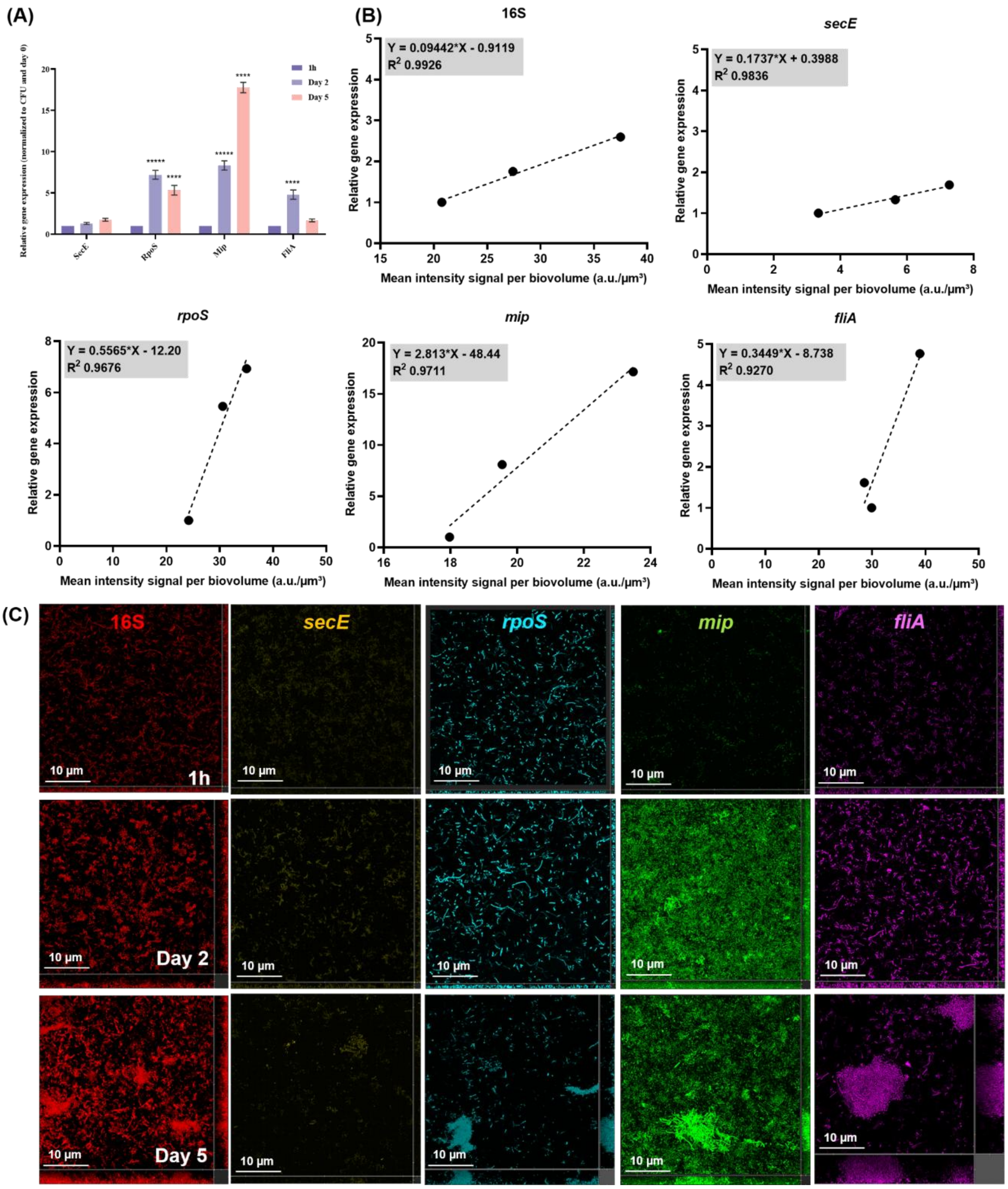


**Supplementary Figure 8 | Quantification and spatial validation of gene expression in *Legionella pneumophila* biofilms.** **(A)** Expression of *secE*, *rpoS*, *mip*, and *fliA* were measured using the *Pfaffl* method after 1 h of initial attachment and on days two and five. Error bars represent standard deviation. * Significant difference between 1 h and the other time points for each gene (*$p$-value < 0.05, **$p$-value < 0.01, ***$p$-value < 0.001, and ****$p$-value < 0.0001). **(B)** Scatter plots showing the relationship between *in situ* singleplex mRNA PNA-FISH signal intensity per biovolume and bulk RT-qPCR measurements for 16S, *secE*, *rpoS*, *mip*, and *fliA* genes after 1 h of initial attachment and on days two and five. The coefficient of determination ($R^2$) is indicated for each regression. **(C)** Confocal microscopy images for the detection of target transcripts at 60 °C, after 1 h of initial attachment and on days two and five under flow conditions.

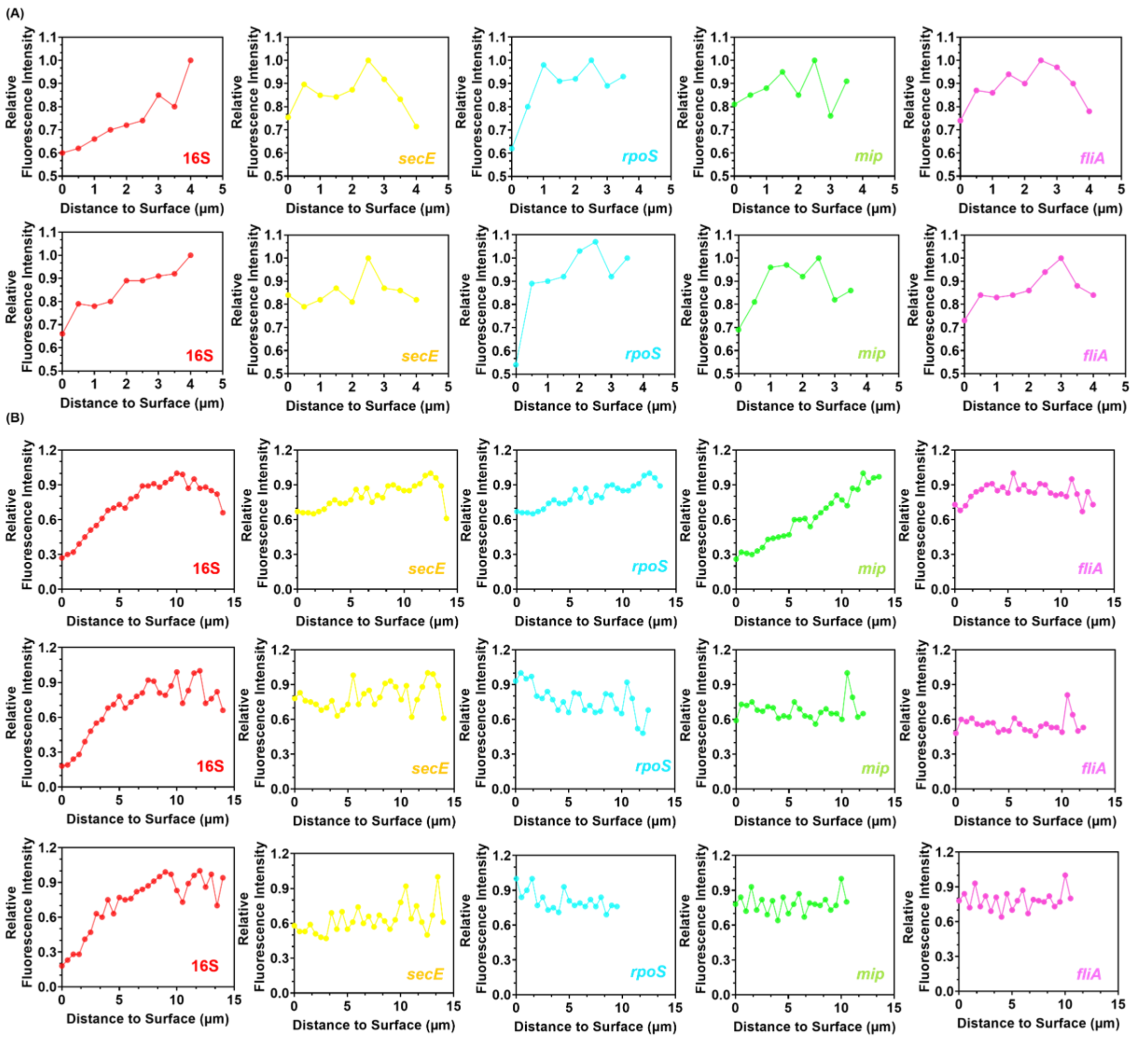


**Supplementary Figure 9 | Spatial organization during biofilm formation.** Vertical spatial profiling of transcript abundance at **(A)** day two for two different microcolonies and at **(B)** day five for three different clusters for 16S, *secE*, *rpoS*, *mip,* and *fliA*.

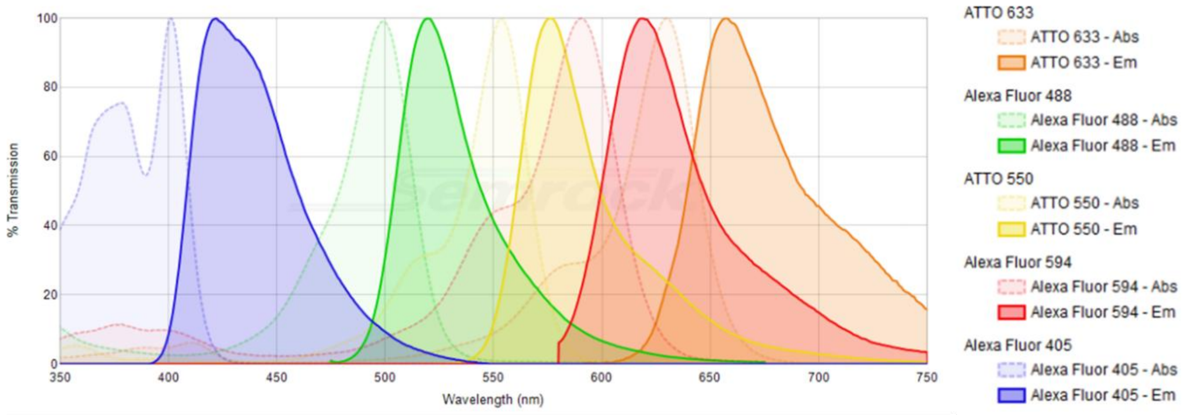


**Supplementary Figure 10 | Spectral profiles of the 5-fluorophore multiplex panel.** Theoretical absorption (dashed lines) and emission (solid lines) spectra of the fluorophores used in this study: ATTO 405, Alexa Fluor 488, ATTO 550, Alexa Fluor 594, and ATTO 633.

**Supplementary Table 1**. Brief description of the gene selection and the characteristics of the PNA probe design.

| Target | Function | Growth phase | Sequence (5'-3') | Length (nt) | Tm (°C) | GC (%) | Gibbs free energy ($\Delta G°$) | Fluorochrome |
|---|---|---|---|---|---|---|---|---|
| **16S** | 16S ribosomal RNA | Exponential/ Transmissive | CTG TAT CGG CCA T | 13 | 70.78 | 54.00 | -14.96 | Alexa Fluor 594 |
| ***secE*** | Translocation of proteins across the cytoplasmic membrane | Exponential phase | CGT GTA GGC CAA AC | 14 | 79.13 | 57.14 | -16.94 | ATTO 405 |
| ***rpoS*** | Stationary-phase sigma factor involved in the transmission phenotype | Exponential/ Transmissive | GCA AGA GTC GCA TA | 14 | 77.19 | 50.00 | -16.21 | ATTO 550 |
| ***fliA*** | Flagellar biosynthesis pathway | Transmissive | GAG GGA GAC TAT CAA | 15 | 80.93 | 46.67 | -16.20 | ATTO 633 |
| ***mip*** | Intracellular replication in protozoa and human macrophages | Transmissive | CCA CTT GGC AAT ACA | 15 | 75.46 | 47.00 | -16.91 | Alexa Fluor 488 |
